# *MATRIN3* deficiency triggers autoinflammation via cGAS-STING activation

**DOI:** 10.1101/2024.04.01.587645

**Authors:** Zohirul Islam, Ahsan Polash, Masataka Suzawa, Bryan Chim, Skyler Kuhn, Sabrina Sultana, Nicholas Cutrona, Patrick T. Smith, Juraj Kabat, Sundar Ganesan, Amir Foroushani, Markus Hafner, Stefan A. Muljo

## Abstract

Interferon-stimulated genes (ISGs) comprise a program of immune effectors important for host immune defense. When uncontrolled, ISGs play a central role in interferonopathies and other inflammatory diseases. The mechanisms responsible for turning on ISGs are not completely known. By investigating MATRIN3 (MATR3), a nuclear RNA-binding protein mutated in familial ALS, we found that perturbing MATR3 results in elevated expression of ISGs. Using an integrative approach, we elucidate a pathway that leads to activation of cGAS-STING. This outlines a plausible mechanism for pathogenesis in a subset of ALS, and suggests new diagnostic and therapeutic approaches for this fatal disease.

**One-Sentence Summary:** Mis-splicing of *Tudor Domain Containing 3* (*TDRD3*) in the absence of MATR3 unleashes R-loops and interferon-stimulated genes.

## Main Text

MATRIN3 (MATR3) is a nuclear RNA-binding protein (RBP) that has diverse functions (*1-8*). However, a better understanding of the link between MATR3 function and human disease is still needed. For instance, mutations in *MATR3* (fig. S1A) have been found in familial amyotrophic lateral sclerosis (ALS) and frontotemporal dementia (FTD) but, mechanistically, how any of them lead to disease pathogenesis is not understood (*9, 10*). Recently, a genetically-engineered mouse model revealed that one of the most common germline missense mutations in MATR3, Serine^85^ to Cysteine (S85C) resulted in neuroinflammation (*11*). However, it is not known whether the effect of MATR3 mutation is cell autonomous, and if so, how such an immune response would be activated. This new paradigm about ALS pathogenesis motivated us to elucidate how perturbing MATR3 can trigger sterile inflammation and identify the key players in this pathway.

Thus, we modelled MATR3 loss-of-function in the human haploid cell line HAP1 (fig. S1, A to D) and performed RNA sequencing (RNA-seq) to identify effects on gene expression (data S1 and S2). Both short- and long-read RNA-seq confirmed that the *MATR3* gene is not expressed in the knock-out (KO) clones (fig. S1B). Loss of MATR3 protein was confirmed by Western blot (fig. S1C), and immunostaining (fig. S1D). This unbiased approach revealed that the *MATR3* KO clones expressed elevated levels of several interferon-stimulated genes (ISGs) in comparison to wildtype (WT) controls (Fig. 1A; data S1 and S2). To validate, we employed Nanostring (data S3), a digital RNA counting technology, and indeed confirmed that several ISG transcripts are increased in *MATR3* KO clones (Fig. 1B). This finding is consistent with the idea that MATR3 deficiency results in a cell autonomous inflammatory response.

**Fig. 1.**
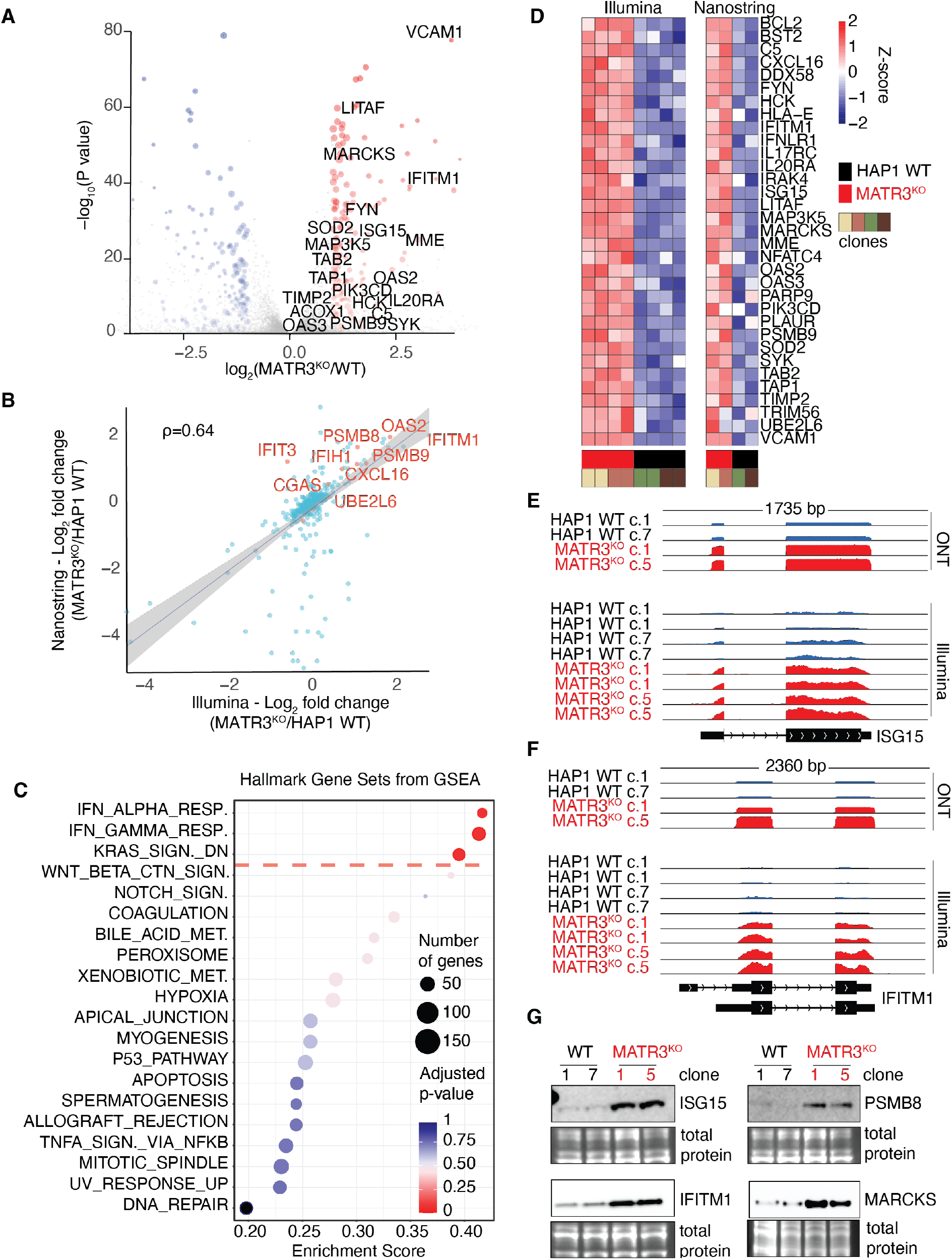
Interferon stimulated genes (ISGs) are elevated in the absence of MATR3. (**A**) Volcano plot depicts differentially expressed transcripts from analysis of Illumina-based RNA-seq of MATR3-deficient HAP1 cells versus WT controls. Each dot represents one transcript. Blue or red indicates transcripts decreased or increased respectively in absence of MATR3. The names of some ISGs are indicated. (**B**) Plot shows correlation between Nanostring and Illumina RNA-seq results. ρ, Spearman’s coefficient. (**C**) Gene Set Enrichment Analysis (GSEA) was performed. Significance and enrichment scores for indicated hallmark gene sets from the Human Molecular Signatures Database (MSigDB) are shown. The hallmark gene sets above the red dashed line have adjusted *P* value < 0.06. (**D)** Heat maps show expression pattern of selected ISGs based on RNA-seq (left) or Nanostring (right). Each column represents the indicated sample. For each transcript, the Z-score of the sample over the row is plotted according to the color scale. (**E, F**) Screenshots of Integrative Genomics Viewer (IGV) depict normalized RNA-seq coverage tracks for the indicated samples either using Oxford Nanopore Technology (ONT) or Illumina. The window length and chromosomal location are indicated above the tracks, while the *ISG15* and *IFITM1* transcript direction (arrowheads) and exon-intron structure based on annotated Reference Sequence (RefSeq) transcripts are indicated below the tracks. (**G)** *Left panels*, Western blots depict abundance of ISG15 and IFITM1 proteins in two *MATR3* KO clones and two WT controls. *Right panels*, same as left but for PSMB8 and MARCKS. Below each blot, an inset of the total protein stain is included as a loading control.

ISGs encode a set of immune system effectors that are part of an important gene expression program which evolved to help the host fight off infections (*12, 13*). When ISGs are aberrantly activated, they are associated with a group of autoinflammatory diseases referred to as interferonopathies (*14*), including Aicardi-Goutières syndrome (AGS). Similar to ALS, a symptom of AGS is sterile neuroinflammation. Gene set enrichment analysis (GSEA) of our RNA-seq data indicated that the interferon (IFN) response is the most significant signature that correlated with MATR3-deficiency (Fig. 1C); this is reminiscent of an interferonopathy. Nanostring validated this IFN signature (Fig. 1D). As two examples of well-known ISGs, we highlight *ISG15* and *Interferon-Induced Transmembrane Protein 1* (*IFITM1*). Both transcripts (Fig. 1, E and F) and proteins (Fig. 1G) were elevated when MATR3 is absent. We also validated a similar pattern of expression for PSMB8 and MARCKS encoded by two additional ISGs (Fig. 1G). To begin mapping the pathway activated by MATR3 deficiency, we will henceforth focus on IFITM1 as a prototypical marker of this inflammatory phenotype.

A well accepted function of MATR3 is that it binds RNA (*5, 7, 15*). However, it is not immediately clear which of the thousands of transcripts reported to be bound by MATR3 would be involved in preventing an innate immune response. To investigate MATR3’s RNA-binding role in our system, we performed photoactivatable ribonucleoside-enhanced crosslinking and immunoprecipitation (PAR-CLIP) to identify transcripts directly bound by endogenous MATR3 (fig. S2A, data S4 and S5). Data from three replicates were highly correlated to each other (Fig. 2A) and revealed on average ∼13 binding sites per transcript from >8,000 genes (fig. S2B). Most of the MATR3 binding sites mapped to introns (Fig. 2B, fig. S2C) which is consistent with its nuclear localization (fig. S1D). Motif analysis using the HOMER algorithm (*16*), or pentamer (5-mer) enrichment found MATR3 binding sites to be pyrimidine-rich (Fig. 2C, fig. S2D). These sequence motifs are consistent with findings from previously published CLIP studies of MATR3 in other cell types (*5, 7, 15, 17*). Next, we checked whether there was a correlation between MATR3 binding and mRNA expression changes in KO versus WT cells. Transcripts bound by MATR3 tended to be down-regulated in KO cells (Fig. 2D), and we subsequently focused our attention on those targets. We hypothesized that the expression of some of these targets might be affected by aberrant splicing.

**Fig. 2.**
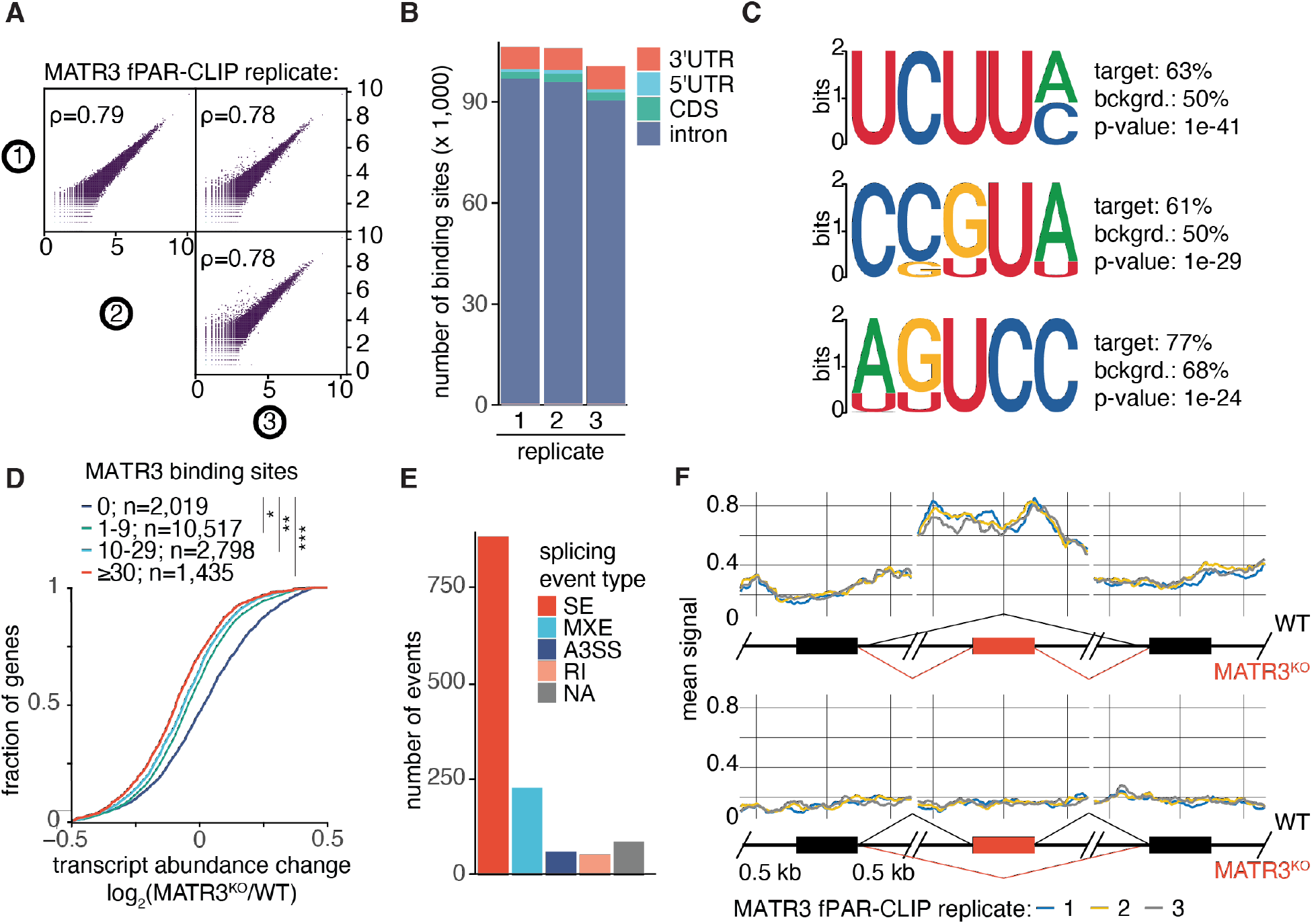
Expression and splicing of transcripts directly bound by MATR3 are affected by loss of MATR3. (**A**) Plots show correlations of crosslinked sequence read frequency (measured by number of T-to-C mutations) on target transcripts for three independent PAR-CLIP replicates (denoted by encircled 1, 2 and 3). ρ, Spearman’s coefficient. (**B**) The fraction of MATR3 binding sites (clusters) are indicated for each annotation category across three independent PAR-CLIP replicates. (**C**) The top three 5-mer sequence motifs identified by HOMER as most significantly enriched among MATR3 binding sites are shown. The nucleotide probabilities for each position are proportional to the height of each character (plotted in bits). Also shown are HOMER enrichment statistics including *P*-values and relative % presence in target versus background (bckgrd.). (**D**) The cumulative distribution of log-transformed fold changes of mRNA abundance comparing MATR3 KO versus WT HAP1 cells as calculated from RNA-seq. MATR3-bound mRNAs were binned based on the number of PAR-CLIP binding sites (clusters): no MATR3 binding sites (blue line); one to nine MATR3 binding sites (green line); 10 to 29 MATR3 binding sites (cyan line); ≥30 MATR3 binding sites (red line). The number of transcripts in each bin is indicated. The statistical significance of the shift compared with unbound mRNAs was determined using a two-sided Kolmogorov-Smirnov (KS) test. *, *P* < 5e-42; **, *P* < 6e-51; ***, *P* < 2.3e-53. (**E**) Bar plot summarizes the number of differential splicing events in each category when comparing MATR3 KO vs WT. Skipped exon (SE), retained intron (RI), mutually exclusive exons (MXE), alternative 5’ splice site (A5SS), and alternative 3’ splice site (A3SS). (**F**) *Top panels*, plots depict MATR3 binding (arbitrary relative units) across region ± 0.5 kb surrounding center of exon included in KO compared to exon upstream or downstream. *Bottom panels*, plots depict MATR3 binding ± 0.5 kb surrounding center of exon excluded in KO compared to exon upstream or downstream.

Previously, MATR3 has been implicated in RNA splicing (*7, 15*). Thus, we employed replicative multivariate analysis of transcript splicing (rMATS) (*18*) to systematically identify alterations in splicing when MATR3 is absent (data S6). This analysis revealed that skipped exons (SE) is the most frequent category of alternative splicing which includes both exon inclusions as well as exclusions (Fig. 2E). MATR3 binding is enriched around exons included in KO (Fig. 2F), but no such enrichment was found at excluded exons suggesting that the latter category is indirectly affected. To prioritize candidate targets for further study, we identified the most down-regulated transcripts in KO which were directly bound by MATR3 and had at least one exon included in KO (data S7). Among this select list, RNA-seq revealed that *Tudor Domain Containing 3* (*TDRD3*) was the most down-regulated transcript in *MATR3* KO cells (other than *MATR3* itself). Since TDRD3 was previously implicated in R-loop metabolism via Topoisomerase III Beta (TOP3B) (*19*) and/or DExH-box helicase 9 (DHX9) (*20*), it became a prime candidate in our investigation.

In the absence of MATR3, exon 5 of *TDRD3* underwent splicing to a previously unannotated, erroneous exon (EE) not apparent in WT (Fig. 3A). The sequence of this EE including the upstream splicing acceptor and downstream donor dinucleotides are evolutionarily conserved among 20 primate species (fig. S3A). Inclusion of this 53 nucleotides-long EE is predicted to disrupt the downstream open reading frame, and results in a premature termination codon (PTC) within exon 8 (fig. S3B) which would trigger nonsense-mediated decay (NMD) thus impeding TDRD3 expression.

**Fig. 3.**
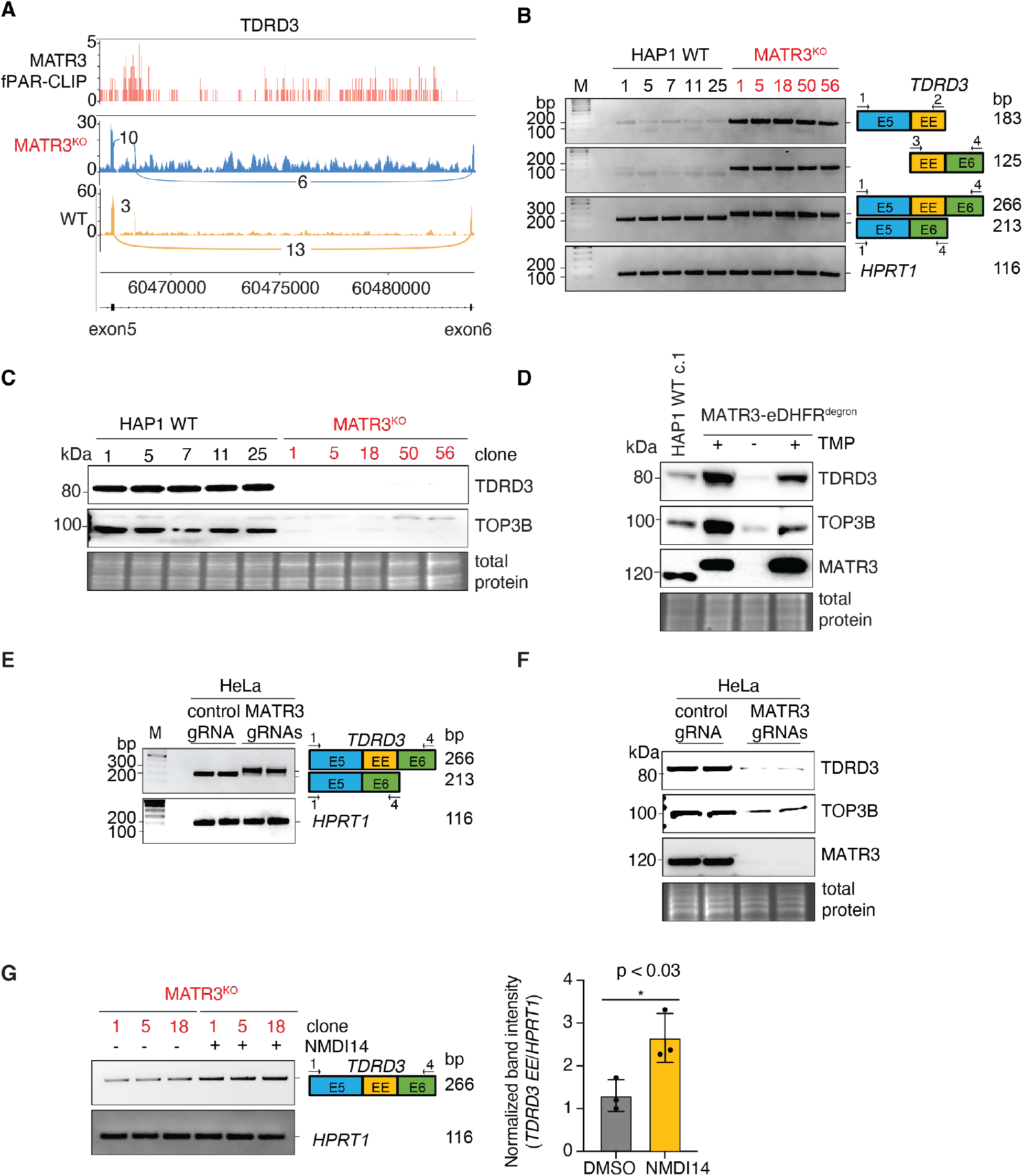
MATR3 is essential for proper splicing and expression of *TDRD3*. (**A**) The top track shows PAR-CLIP sequence read coverage representing endogenous MATR3 binding in WT HAP1 cells (pink). The Sashimi plots depict the RNA-seq coverage for two biological replicates of MATR3 KO (blue) and WT controls (yellow) for a region starting from exon 5 of *TDRD3* (transcribed from left to right) and ending at exon 6. An unannotated erroneous exon (near exon 5) is included in MATR3 KO. The arcs represent splicing, and the numbers of supporting splice junction reads are indicated for each. (**B**) RT-PCRs of total RNA from five *MATR3* KO clones and five WT controls were performed to assay differential *TDRD3* exon usage and *HPRT1* mRNA as a housekeeping control. Following agarose gel electrophoresis, images depict: *top panel*, 183 bp RT-PCR product representing *TDRD3* isoform in which exon 5 is spliced to erroneous exon (EE); *second panel, TDRD3* isoform in which EE is spliced to exon 6 (125 bp); *third panel*, canonical *TDRD3* isoform in which exon 5 is spliced to exon 6 (213 bp), as well as the aberrant *TDRD3* isoform in which EE is included in between exons 5 and 6 (266 bp). (**C**) Western blots depict abundance of TDRD3 (*top*) and TOP3B (*middle*) proteins in four *MATR3* KO clones and four WT controls. *Bottom*, an inset of the total protein stain is included as a loading control. (**D**) HAP1 cells engineered to express endogenous MATR3 that is eDHFR-tagged were grown in 10 μM trimethoprim (TMP) to stabilize MATR3-eDHFR fusion protein (lane 2); then washed and grown in TMP free (-TMP) media for 96 h to deplete MATR3-eDHFR (lane 3); and then in 10 μM TMP for 48 h to restore MATR3-eDHFR expression (lane 4). Western blots of TDRD3 (*top*), TOP3B (*second panel*) and MATR3 (*third panel*) are shown. *Bottom*, an inset of the total protein stain is included as a loading control. (**E**) A pool of HeLa cells nucleofected with recombinant Cas9 protein complexed with either control gRNA or a pair of gRNAs targeting *MATR3* were harvested for analysis 72 hours after nucleofection. RT-PCRs were performed to measure abundance of mis-spliced *TDRD3* transcript and *HPRT1* mRNA as a housekeeping control. Following agarose gel electrophoresis, images depict: *top panel*, canonical *TDRD3* isoform in which exon 5 is spliced to exon 6 (213 bp), as well as the aberrant *TDRD3* isoform in which EE is included in between exons 5 and 6 (266 bp). (**F**) Samples are same as in E. Representative Western blots of TDRD3 (top), TOP3B (second panel) and MATR3 (third panel) are shown. *Bottom*, an inset of the total protein stain is included as a loading control. (**G**) RT-PCRs of total RNA from three *MATR3* KO clones treated with either the NMD inhibitor, NMDI14, or DMSO solvent-only control were performed to measure relative abundance of mis-spliced *TDRD3* transcript and *HPRT1* mRNA as a housekeeping control. Following agarose gel electrophoresis, images depict: *top panel*, the aberrant *TDRD3* isoform in which EE is included in between exons 5 and 6 (266 bp). Band densities were quantified and plotted in graph (right panel).

To address the potential of clonal variability, we generated three additional *MATR3* KO clones and isolated three matching WT clones to strengthen our validation studies going forward (fig. S4, A and B). This novel *TDRD3* mis-splicing event could be validated by three different assays employing reverse transcription followed by polymerase chain reaction (RT-PCR) in five independent *MATR3* KO clones (Fig. 3B). In contrast, the mis-splicing is not detected in WT controls (Fig. 3B). Ultimately, not only is *TDRD3* mRNA reduced by this EE inclusion (data S1), expression of TDRD3 protein is severely hampered in the absence of MATR3 (Fig. 3C). TDRD3 forms a heterodimer with TOP3B protein (*19, 21*), whose expression is also affected by MATR3 deficiency (Fig. 3C). To validate our findings using an orthogonal approach (fig. S5A), we tagged endogenous MATR3 with a “degron” derived from *E. coli* dihydrofolate reductase (eDHFR) which facilitates reversible drug-inducible targeted protein degradation (*22*). Indeed, acute depletion of MATR3 protein for 96 hours (fig. S5B), also reduced expression of both TDRD3 and TOP3B (Fig. 3D). Furthermore, restoring MATR3 protein expression for 48 hours, rescued both defects which supports the idea that MATR3 directly regulates TDRD3 expression and indirectly TOP3B protein stability. As evidence that our findings will translate to other cell types, deletion of *MATR3* in HeLa, a human cervical carcinoma cell line, also resulted in mis-splicing of TDRD3 (Fig. 3E) and reduced protein expression of TDRD3 and TOP3B (Fig. 3F). Finally, treatment of MATR3 KO cells with NMDI-14, a small molecule inhibitor of NMD (*23*), increased expression of this aberrant *TDRD3* transcript (Fig. 3G). This result supports the idea that this mis-spliced *TDRD3* transcript is degraded via NMD.

Since TDRD3 and TOP3B have been implicated in the metabolism of R-loops (*19, 20*), we visualized RNA-DNA hybrids in MATR3 KO cells using the S9.6 monoclonal antibody. We found aberrant accumulation of RNA-DNA hybrids in the cytoplasm of MATR3 KO cells compared to WT controls (Fig. 4A). The immunostaining signal is degraded by Ribonuclease H (RNaseH) treatment which supports the notion that they represent RNA-DNA hybrids. We hypothesize that these cytoplasmic RNA-DNA hybrids serve as the trigger to induce expression of ISGs such as *IFITM1* (Fig. 4B). Using the reversible degron system, we demonstrate that IFITM1 expression is induced by acute MATR3 depletion, and inhibited when MATR3 is restored (Fig. 4C). Previously, it was demonstrated that RNA-DNA hybrids can be sensed by Cyclic GMP-AMP Synthase (cGAS), resulting in catalysis of 2’,3’-cyclic GMP-AMP (cGAMP) which in turn activates Stimulator of Interferon Response cGAMP Interactor 1 (STING1) (*24, 25*). To determine whether activation of the cGAS-STING pathway is responsible for the autoinflammatory response seen in *MATR3* KO cells, we deleted *CGAS*. Indeed, loss of cGAS abrogated induction of IFITM1 in the absence of MATR3 (Fig. 4D). To demonstrate that TDRD3 was critical for preventing induction of IFITM1, we generated *TDRD3* KO clones. Consistent with our working model (fig. S6), in the absence of TDRD3, expression of TOP3B protein is reduced, and as a result, IFITM1 is induced (Fig. 4E). Finally, to prove that the erroneous exon (EE) in *TDRD3* is causing the central defect in MATR3-deficiency, we deleted that EE in *MATR3* KO clones. Remarkably, simply removing the EE fully rescued TDRD3 protein expression in *MATR3* KO cells (Fig. 4F).

**Fig. 4.**
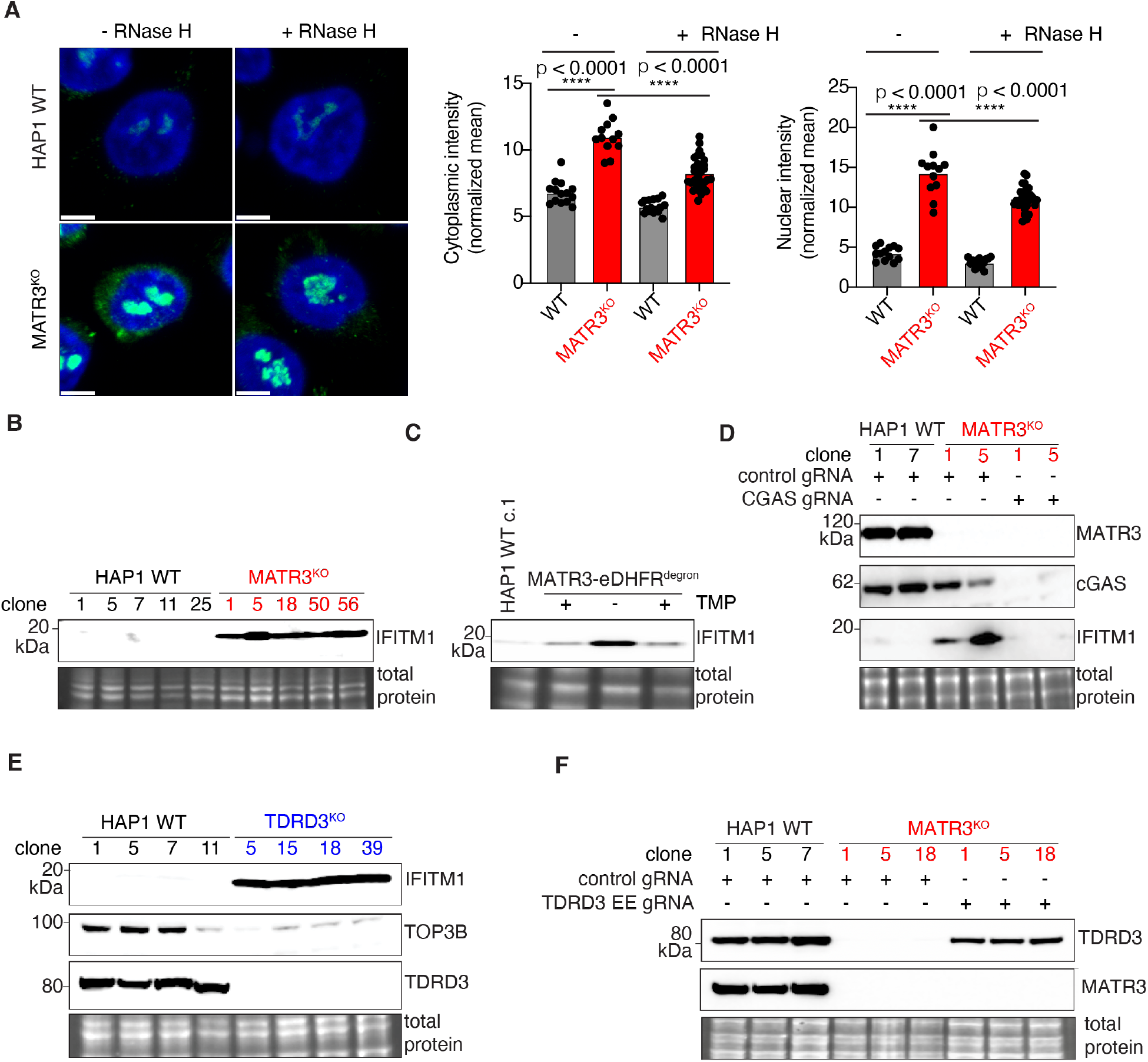
An innate immune response starts from MATR3, then TDRD3, TOP3B, cGAS and ultimately IFITM1. (**A**) Representative images show S9.6 immunostaining (green) which detects RNA-DNA hybrids in MATR3 KO vs WT. RnaseH treatment to digest RNA-DNA hybrids was done to demonstrate specificity. DAPI (blue) was used to stain nuclei. The fluorescence intensity in either cytoplasm (middle panel) or nucleus (right panel) of single cells from indicated samples are plotted in graphs. (**B**) Representative Western blot depicts abundance of IFITM1 in five *MATR3* KO clones and five WT controls. *Bottom*, an inset of the total protein stain is included as a loading control. (**C**) Representative Western blots of IFITM1 (*top*), and MATR3 (*middle*) in a pool of HAP1 cells engineered to express endogenous MATR3 that is eDHFR-tagged. Cells were grown in 10 μM TMP to stabilize MATR3-eDHFR fusion protein (lane 2); then washed and grown in TMP free (-TMP) media for 96 h to deplete MATR3-eDHFR (lane 3); and then in 10 μM TMP for 48 h to restore MATR3-eDHFR expression (lane 4). *Bottom*, an inset of the total protein stain is included as a loading control. (**D**) Representative Western blots of MATR3 (top), cGAS (second panel) and IFITM1 (third panel) are shown. Two independent replicates of WT HAP1 cells were nucleofected with Cas9 complexed with control gRNA. Two independent replicates of *MATR3* KO cells were nucleofected with Cas9 complexed with either control gRNA or a pair of gRNAs targeting *CGAS*. Cells were harvested 72 hours after nucleofection. *Bottom*, an inset of the total protein stain is included as a loading control. (**E**) Representative Western blots depict abundance of IFITM1 (*top*), TOP3B (*second panel*) and TDRD3 (*third panel*) proteins in four *TDRD3* KO clones and four WT controls. *Bottom*, an inset of the total protein stain is included as a loading control. (**F**) Western blots of TDRD3 (top) and MATR3 (second panel) are shown. Three independent replicates of WT HAP1 cells were nucleofected with Cas9 complexed with control gRNA. Three independent replicates of *MATR3* KO cells were nucleofected with Cas9 complexed with either control gRNA or a pair of gRNAs targeting EE of *TDRD3*. Cells were harvested 72 hours after nucleofection. *Bottom*, an inset of the total protein stain is included as a loading control.

In summary, by identifying HAP1 cells as a useful model system to study MATR3 loss of function, we discovered a previously unappreciated pathway that can trigger the expression of ISGs in a cell autonomous fashion. It appears that one important function of MATR3 is to prevent mis-splicing of *TDRD3* and the subsequent activation of ISGs. In the absence of MATR3, a novel EE within *TDRD3* is included following its transcription and splicing, and results in a premature termination codon (PTC), a trigger for NMD. Ultimately, TDRD3 protein levels are markedly reduced along with its heterodimeric partner, TOP3B. This leads to aberrant accumulation of RNA-DNA hybrids in the cytoplasm of MATR3-deficient cells which activates the cGAS-STING pathway and ultimately ISGs. These findings imply a potential pathogenetic mechanism that should be considered in the setting of ALS, particularly those with *MATR3* mutations. In addition, our results nominate defects in TDRD3 and TOP3B as potentially unappreciated causes of ALS and interferonopathies. Similar to the mechanism we uncovered, one frequent cause of AGS is the mutation of Ribonuclease H2 (*26*), an enzyme that is well known for digesting RNA-DNA hybrids within cells (*27*). Our results suggest a range of options that might aid in the diagnosis of ALS, and perhaps even its treatment. For example, in ALS cases where neuroinflammation is an early event, pharmacological targeting of the cGAS-STING pathway might inhibit disease progression. Finally, since R-loop metabolism is implicated in tumorigenesis (*28*), the pathway we highlight here may also be linked to outcomes of certain cancers.

## Acknowledgments

We thank Gustavo Gutierrez-Cruz, Faiza Naz and Stefania Dell’Orso (Genomic Technology Section, NIAMS) for Illumina sequencing; the NIAID Office of Cyber Infrastructure and Computational Biology for high-performance computing; Pedro Batista (National Cancer Institute), Astrid Haase (National Institute of Diabetes and Digestive and Kidney Diseases), Chrysi Kanellopoulou (Incyte Research Institute), Philipp Oberdoerffer (Johns Hopkins University) and members of S.A.M.’s group for comments and suggestions. This research was supported by the Division of Intramural Research of NIAID/NIH and NIAMS/NIH.

## Funding

Division of Intramural Research of NIAID/NIH ZIA AI001185 (SAM) Division of Intramural Research of NIAMS/NIH ZIA AR041205 (MH)

## Author contributions

Conceptualization: MH, SAM Methodology: ZI, AP, MS, BC, SK, JK

Investigation: ZI, AP, MS, BC, SK, SS, NC, PTS, SG, AF

Visualization: ZI, AP, BC, MH, SAM

Funding acquisition: MH, SAM

Project administration: SAM Supervision: MH, SAM

Writing – original draft: SAM

Writing – review & editing: ZI, MH, SAM

## Competing interests

Authors declare that they have no competing interests.

## Data and materials availability

Almost all data are available in the main text or the supplementary materials. The RNA-seq (GSE216825 and GSE216826), Nanostring (GSE216829) and PAR-CLIP data (GSE262647) have been deposited in The NCBI Gene Expression Omnibus (https://www.ncbi.nlm.nih.gov/geo/) under the indicated accession numbers.

## Supplementary Materials

Materials and Methods

Figs. S1 to S6

Tables S1 to S2

References (*1*–*20*)

Movies S1

Data S1 to S7

## Notes

### Competing Interest Statement

The authors have declared no competing interest.

