## Supplement for "*MATRIN3* deficiency triggers autoinflammation via cGAS-STING activation"

### Materials and Methods

Cell culture and treatments

HAP1 cells were cultured in Iscove's Modified Dulbecco's Medium (IMDM) (Quality Biological, cat#112-035-101) supplemented with 10% FBS (Gemini, cat#100-106) and 1% Pen/Strep (Gibco, cat#15140-122). MATR3 KO clone 1 (c.1) and clone 5 (c.5) were purchased from Horizon Discovery (cat# HZGHC007500c001 and cat# HZGHC007500c005) along with wildtype controls (cat# C631_29281 batch 40661, cat# C631_29281, batch 50047 later mentioned as HAP1 WT clone 1 (c.1) and clone 7 (c.7), respectively. HeLa cells were cultured in Dulbecco's modified Eagle Medium (DMEM) (Gibco, cat# 10569-010) supplemented with 10% FBS (Gemini, cat# 100-106) and 1% Pen/Strep (Gibco, cat# 15140-122). To generate knockout (KO) cell lines, parental cells were nucleofected with recombinant Cas9 complexed with CRISPR guide RNAs (gRNAs) comprised of crRNA:tracrRNA (IDT, cat# 1072533) duplexes. Alt-R® S.p. Cas9 Nuclease V3 (IDT, cat# 1081058), Alt-R® CRISPR crRNAs (IDT, see table S1) and electroporation enhancer (IDT, cat# 1075915) were used per manufacturer's recommendations. For HAP1 and HeLa cells, the SE cell line 4D-Nucleofector kit S (Lonza) were used with a 4D nucleofection unit (Lonza) using protocol DZ-113. Briefly, $8x10^5$ cells were centrifuged at 300xg for 5 min at room temperature. Supernatant was removed and cells resuspended in 20 µl of Nucleofector solution SF, and transferred into an Rnase-, Dnase-free tube. RNP complex was prepared by combining Alt-R guide RNA 1.2 µl (120 pmol from 100 pmol/mL stock), Alt-R Cas9 1.7 µL (104 pmol from 61µM stock), and 2.1 µL PBS in sterile and Rnase-, Dnase-free tube and incubated at room temperature for 20 minutes. Next, 5 µL of the RNP complex was added to the cell suspension. Finally, 1 µL of Alt-R Cas9 Electroporation Enhancer was added to each tube. After mixing, 26 µL of the cell:RNP complex mixture were transferred to a well of the 16-well Nucleocuvette module. Bubbles, if present, were removed by gently tapping the Nucleocuvette module. After electroporation, cells were resuspended in culture medium and incubated in a 37° C, 5% $CO_2$ incubator. Single cell clones were isolated using limiting dilution and were screened by PCR genotyping assays (see table S1 for primers). The PCR products were also submitted for Sanger sequencing (Psomagen) to determine the exact deletion or insertion in each KO clone. Several WT clones were kept as matching controls and validated by Western. When indicated, the pool is analyzed directly 72 h after nucleofection.

To inhibit nonsense-mediated RNA decay (NMD), cells were treated with 5µM NMDI14 (MCE, cat# HY-111374) or DMSO for 6 hrs.

Degron

To deplete endogenous MATR3 protein in an inducible manner, we tagged its C-terminus with an optimized *E. coli* dihydrofolate reductase (eDHFR) degron which is regulated by trimethoprim (TMP), its small-molecule ligand (*1*). The eDHFR-tagged fusion protein is degraded by the proteosome in the absence of TMP, but stabilized when it binds to TMP. PCR primers used to generate the homology-directed repair (HDR) donor are listed in the Supplementary Material (table S1). Two phosphorothioate linkages were included at the 5′ end of each primer to prevent degradation of the HDR donor by cellular exonucleases. The plasmid pAc5 HA3-eDHFR-T2A-puro was a gift from David Bentley (Addgene plasmid # 86395;

http://n2t.net/addgene:86395). It was used as template for PCR using Phusion® High-Fidelity DNA Polymerase per manufacturer's instructions (NEB, cat# M0531S). We used CRISPR-mediated homologous recombination essentially as above to insert the cassette in place of MATR3's STOP codon (exon 14). After nucleofection, the cells were selected using 10 μg/mL of puromycin (Gibco, cat# A11138-03) to select for successful integration of HA3-eDHFR-T2A-puro cassette. 10 μM TMP was added to prevent degradation of MATR3-eDHFR fusion protein. When the cells look healthy after 1–2 weeks of puromycin selection, genomic DNA was isolated for PCR genotyping. The primers for genotyping are listed in table S1. Expression of MATR3 protein ± TMP was validated by Western.

Western blotting

70 to 80% confluent cells were harvested with a lysis buffer: 20mM Tris (pH 7.5), 150mM KCl, 3mM $MgCl_2$, 0.3% NP-40 and 10% glycerol) supplemented with PhosSTOP (Sigma, cat# 04906837001) and cOmplete tablets, mini (Sigma, cat# 04693159001) on ice. The lysate was sonicated two times for 5s at 60% amplitude (Heat Systems-Ultrasonics W-225R Sonicator) to shear chromatin and centrifuged at 14000 rpm for 15 mins, and the supernatant was collected and denatured at 70° C for 10 mins. Protein concentration was measured using BCA method (Pierce BCA protein assay kit). Equal amounts of protein were loaded onto a NuPAGE 4-12% Bis-Tris Gel (Invitrogen) and transferred onto a PVDF membrane (Novex, Life technologies) with a Novex X Cell Western Blot Semidry system (Thermo Fisher Scientific). Blot was stained with TotalStain Q (Azure, cat# AC2225) and imaged to visualize protein transfer and loading. Then, the membrane was blocked with 5% (wt/vol) milk in Tris Buffer Saline with 0.1% Tween (TBST buffer) for 1 h at room temperature with gentle rocking. Next, the membrane was incubated with a primary antibody (table S2) at 4° C overnight with gentle rocking, and the blots were washed for 5 min with TBST three times. Subsequently, an HRP-conjugated secondary antibody incubation was followed for 1 h at room temperature and washed again for 5 min with TBST three times. Detection was performed using the SuperSignal West Dura Extended Duration Substrate (Thermo Fisher Scientific, cat# 34075), and images were acquired using Azure c500 Imager (Azure Biosystems). Band densities on blots were quantified with ImageJ (NIH), and background from an adjacent blank area of the blot was subtracted. The signal from the protein band of interest was normalized to total protein stain.

Immunofluorescence microscopy

For MATR3 and PTBP1 immunostaining, cells were seeded in μ-slide 8-well (ibidi, cat# 80826). After 24 h, cells were washed 3 times with 1x PBS (Gibco, cat#10010023). Next, cells were fixed with 4% paraformaldehyde (PFA) (Electron microscopy Sciences, cat# 15170) and permeabilized with 0.1% TritonX-100. After blocking with 5% BSA for 1 h at room temperature, cells were incubated with primary antibody overnight at 4° C followed by a fluorescent secondary antibody. Nuclei staining was done with 5ng/ml DAPI in 1XPBS for 5 mins at room temperature. Images were taken using Leica SP8 confocal microscope (Research Technologies Branch, NIAID)

For S9.6 immunostaining, cells were washed once with 1× PBS and fixed with ice-cold 100% methanol for 5 min at −20°C. Cells were washed briefly twice with 1× PBS. RNase H (NEB, cat# M0297S) was diluted 1:50 in 1× RNase H buffer and incubated for 6 hours at 37°C. Mock-treated samples were incubated in parallel in the same reaction buffer but without enzymes added. Following incubation, samples were washed twice in 0.5% PBST, then once in

PBS for 5 min each, and blocked with staining buffer (5% BSA in PBS) for 30 min at 37°C. Samples were incubated in staining buffer with a 1:25 dilution of S9.6 antibody overnight at 4°C. After washing thrice with 0.5% PBST for 10 mins, samples were incubated with a 1:1,000 dilution of anti-mouse Alexa Fluor 488–conjugated secondary antibodies for 1 h. Samples were washed thrice for 10 min with 0.5% PBST. DAPI was used as nuclear counterstain. Images were taken using Leica SP8 confocal microscope (Research Technologies Branch, NIAID).

Image quantitation

We used Imaris software package (version 10.1, Andor Technology Inc., Concord, MA) for image processing. We rendered surface objects as region of interest (ROI) specific for nucleoli. This ROI was then used to create an additional channel. The new channel contained all the cell signal voxels from the first channel except for the nucleoli signal. We utilized the Cell module in Imaris to create a cell model containing both the nucleus and cytoplasm. We exported intensities from all channels within the nuclei and cytoplasm and used them to generate graphs for mean intensity comparisons and to perform t-test in Prism (Graphpad Software).

RNA isolation

Total RNA was isolated using RNAzol (Sigma, cat# R4533-100ML) per manufacturer's instruction. RNA was quantified with Qubit RNA BR Assay Kit (Biotium, cat# 31073) and Qubit 4 Fluorimeter according to manufacturer's instruction. RNA integrity was verified with HS RNA Kit (15NT) on the 5200 Fragment Analyzer (Agilent Technologies, cat# DNF-472-500) per manufacturer's protocol.

RT-PCR

First-strand cDNA was synthesized from one µg of total RNA using SuperScript IV First-Strand Synthesis System (Thermo Fisher Scientific, cat# 18091050) and oligo dT primers. Polymerase chain reaction (PCR) was done by using Platinum™ SuperFi II Green PCR Master Mix (ThermoFisher, cat# 12369010). Real-time quantitative PCR was performed using iTaq™ Universal SYBR® Green Supermix (Bio-Rad, cat# 1725038) in a c1000 Touch Thermal Cycler, CFX96 Real-Time System (Bio-Rad) and primers for *TDRD3*, and *HPRT1* are listed on Table S1.

Illumina RNA-seq

Cells were lysed by RNAzol (Sigma, cat# R4533-100ML) to prepare total RNA for sequencing using the DirectZol RNA miniPrep Plus (Zymo research, cat# R2072) per manufacturer's instruction. cDNA libraries were prepared with the NEBNext® rRNA Depletion Kit v2 (NEB, cat# E7405L) followed by NEBNext® Ultra™ II Directional RNA Library Prep Kit for Illumina (NEB, cat# E7760L) and NEBNext® Multiplex Oligos for Illumina (Unique Dual Index UMI Adaptors RNA Set 1, cat# E7416L) per manufacturer's instruction. The final libraries were sequenced on Illumina NovaSeq6000, Chemistry 1.5V as 100-8-8-100 by the Genomic Technology Section (NIAMS, NIH).

Illumina RNA-seq analyses

The raw base calls were demultiplexed and converted into FastQ format using bcl2fastq. The raw FastQ files were processed using version 1.8.0 of the OpenOmics/RNA-seek pipeline (https://github.com/OpenOmics/RNA-seek). This pipeline orchestrates a comprehensive set of data-processing and quality-control steps optimized for RNA-sequencing data. Briefly, the quality of each sample was assessed using FastQC v0.11.9 (https://www.bioinformatics.babraham.ac.uk/projects/fastqc/), Preseq v2.0.3 (*2*), Picard tools v2.17.11 (https://broadinstitute.github.io/picard/), FastQ Screen v0.9.3 (*3*), Kraken2 v2.0.8 (*4*), QualiMap (*5*), and RSeQC v2.6.4 (*6*). Adapter sequences were trimmed using Cutadapt v1.18 (*7*). The trimmed reads were aligned GENCODE's (*8*) human primary genome assembly, GRCh38.p13, using the splice-aware aligner STAR version 2.7.6a (*9*). Gene and transcript expression levels were estimated with GENCODE's comprehensive human gene annotation (release 42) via RSEM v1.3.3 (*10*). Differentially expressed genes were identified by DESeq2 (*11*). For GSEA analysis, the normalized count obtained from DESeq2 was used as input for the GSEA software (*12, 13*). Alternative splicing events were identified using rMATS v4.1.1 (*14*). Significant differentially expressed alternative splicing events were identified using the following thresholds: "FDR <= 0.05" and "(average(IJC_WT)>=5 or average(IJC_KO)>=5 or average(SJC_WT)>=5 or average(SJC_KO)>=5)" and "abs(IncLevelDifference)>=0.2".

ONT RNA-seq and analyses

The ONT PCR-cDNA sequencing kit (Oxford Nanopore Technologies, cat# SQK PCS-109) was used for library preparation per manufacturer's instructions. For each sample, 50 ng of input total RNA was used. Libraries were sequenced on the (Oxford Nanopore Technologies, cat#FLO-MIN106D) and the MinKNOW software was used to basecall reads and generate fast5 files.Reads were mapped to hg38 using minimap2 (*15*) using the following parameters:- x map-ont -secondary=no -ax splice –uf . The resulting bam files were then filtered for mapping quality and unique mapping using samtools view -F 256 -q 20, and converted to RPM-normalized bigwig files. The filtered bam file were used along with Gencode v 33 as input for Subread's (version 2.0.1) feature count (parameters: -L -t exon -g gene_id) to quantify gene level expression (*16*). Differential expression was determined by DESeq2 version 1.22.2 (*11*).

Nanostring and analyses

The reporter codeset used was the nCounter Human Host Response Panel (cat# XT-HHR-12), which measures the absolute abundance of 785 transcripts associated with host response, including host susceptibility, interferon response, innate immune cell activation, adaptive immune response, homeostasis, and 12 housekeeping genes. Detailed information is provided on their website https://nanostring.com/products/ncounter-assays-panels/immunology/host-response/. Total RNA 100ng from fresh frozen samples was used for mRNA-expression analysis on the nCounter instrument (Nanostring Technologies, Seattle, USA). Target mRNA was hybridized with reporter-capture probes at 65 °C for 24 h, according to the manufacturer's protocol. The probe–target complexes were aligned and immobilized in the nCounter cartridge, which was then placed in the nCounter SPRINT to acquire images and process data, following the manufacturer's protocol. Center for Cancer Research Genomics Core Facility (NCI) performed data acquisition.

Processed RCC files were imported into nSolver Analysis Software v4.0.70. All samples passed the default nSolver QC checks for Imaging, Binding Density, Positive Control Linearity and Positive Control Limit of Detection. Background Subtraction and Thresholding steps were skipped and raw gene expression counts were normalized using the geometric means of the Positive Controls (POS_A, POS_B, POS_C, POS_D, POS_E, POS_F) and Housekeeping genes (ABCF1, ALAS1, GUSB, HPRT1, MRPS7, NMT1, OAZ1, PGK1, SDHA, and TBP). Two housekeeping genes, NRDE2 and STK11IP, were excluded from the normalization due to low average counts (<100). All samples fell within the default Threshold Min and Threshold Max values for Positive Control Normalization (Min 0.3, Max 3.0) and CodeSet Content Normalization (Min 0.1, Max 10.0). Ratio Data analysis was performed with default parameters to compute fold change estimates, t-test P values and 95% confidence intervals between the MATR3 KO and WT samples. Normalized counts, fold changes and P values were imported into RStudio v1.4.1717 (running R v4.1.1) to generate gene expression heatmaps (R package: pheatmap) and volcano plots (R package: ggplot2). The nSolver processed data can be found in data S3.

PAR-CLIP

PAR-CLIP was performed as previously described (*17*) with modifications in the cross-linked cell preparation and immunoprecipitation steps to optimize for MATR3. Briefly, 60 $\times 10^6$ HAP1 WT or MATR3$^{KO}$ cells grown in three 10 cm$^2$ plates were used for each biological replicate. 4-thiouridine (4SU) was added to the cell culture medium at a final concentration of 250 μM for 4 hours. After washing with ice-cold PBS, cells were cross-linked (irradiated with 312 nm UV light, 5 min) and scraped off the dishes by pipetting.

Cell Lysis, RNaseI Treatment: The cross-linked cell pellet was lysed in 3x volume of IP buffer (20mM Tris-HCL pH7.5, 150 mM NaCl, 2mM EDTA, 1.0% v/v NP-40, 0.5mM dithiothreitol) supplemented with cocktail of protease inhibitors (Sigma, cat# 04693159001) and phosphatase inhibitors (Sigma, cat# 4906845001). The lysate was left on ice for 10 min and centrifuged at 13,000g for 15 min at 4 °C. The supernatant was treated with RNaseI (Invitrogen, cat# AM2294) at a final concentration of 0.1U/uL for 10 min at 22 °C, then cooled on ice for 5 min. Immunoprecipitation: 40uL of Dynabeads ProteinA (Invitrogen, cat# 10001D) was mixed with 3ug of anti-MATR3 (Bethyl, cat# A300-591A), incubated for 1 hour at 4°C with rotation, then washed three times with IP buffer. Anti-MATR3 binding beads were added to the RNase-treated supernatants and incubated for 2 hours at 4°C with rotation. The beads were washed three times with 1 ml of IP buffer. The further procedure (RNaseI digestion, dephosphorylation, 3′ adaptor ligation, phosphorylation, SDS PAGE separation, gel excision, Proteinase K digestion, 5′ adaptor ligation, reverse transcription, sequencing library preparation) were performed as described previously (*17*). Sequencing was performed as single reads on an Illumina NextSeq2000 system (Genomic Technology Section, NIAMS).

PAR-CLIP analyses

PAR-CLIP data were analyzed using a PARpipe pipeline that utilized PARAlyzer for peak calling. Within the pipeline, Bowtie1 was used to align the fastq files with the Human reference genome (hg38) using the following parameters: ` -v 2 -m 2 --all --best --strata`. The resulting `clusters.csv` files contained all the metrics for ranking and filtering the clusters. Deep tools were used to compare the similarity of the three replicates of PAR-CLIP. Briefly, deepTools was used to calculate several aligned reads across uniform genomic bins. The normalized reads were used to compute Pearson's correlation coefficient. The consensus motif was calculated using Homer's top 3000 filtered annotated clusters (v4.10). CDF plots were performed to evaluate the relationship of transcriptome stability with PAR-CLIP binding with genes with basemean greater than 100 in differential expression. They belonged to the top 80% for the sum of T to C conversions.

### Figure S1

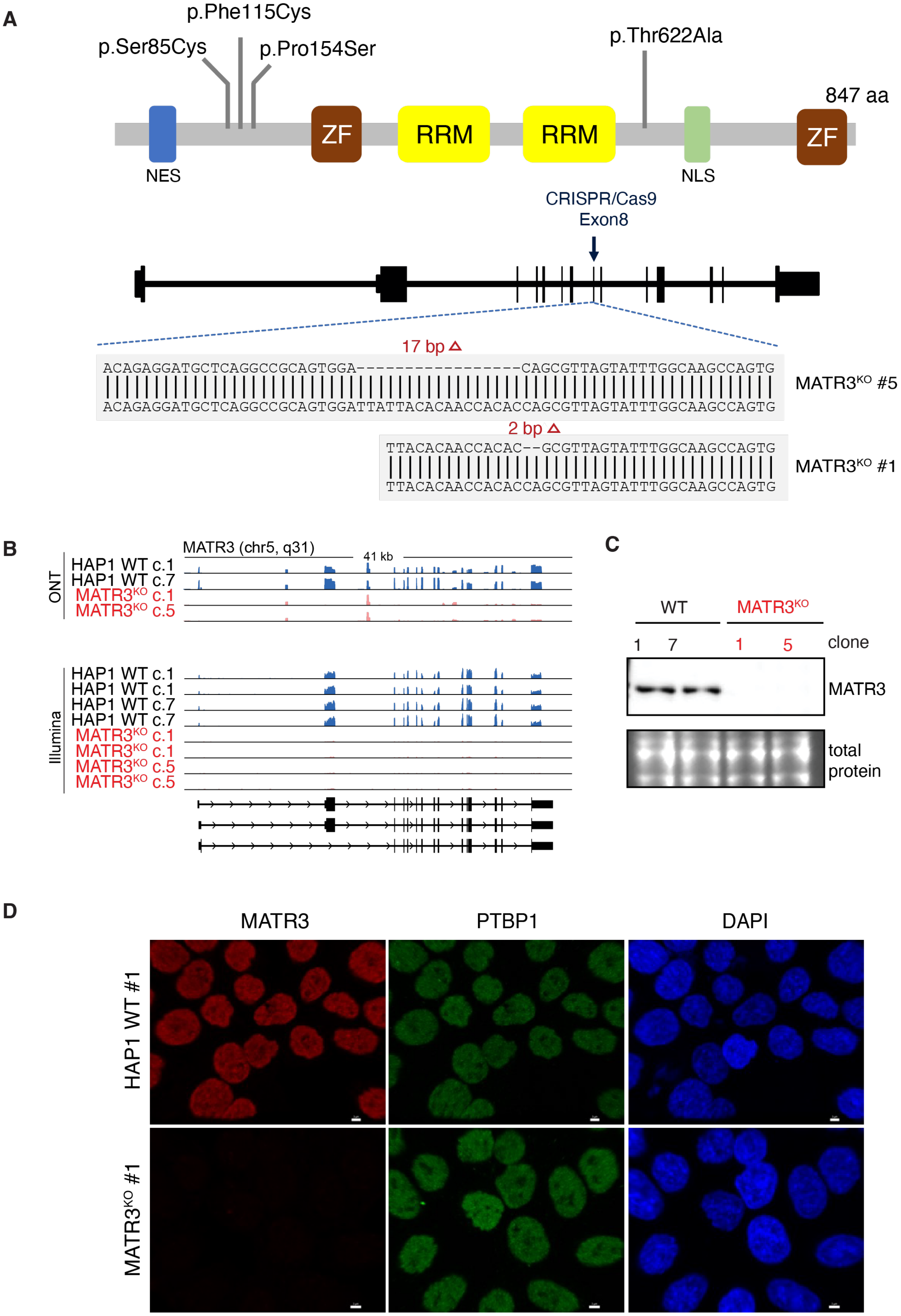

A
p.Phe115Cys
p.Ser85Cys
p.Pro154Ser
p.Thr622Ala
847 aa
ZF
RRM
RRM
ZF
NES
NLS
CRISPR/Cas9
Exon8
17 bp Δ
ACAGAGGATGCTCAGGCCGCAGTGGA-----------------CAGCGTTAGTATTTGGCAAGCCAGTG
ACAGAGGATGCTCAGGCCGCAGTGGATTATTACACAACCACACCAGCGTTAGTATTTGGCAAGCCAGTG
MATR3KO #5
2 bp Δ
TTACACAACCACAC--GCGTTAGTATTTGGCAAGCCAGTG
TTACACAACCACACCAGCGTTAGTATTTGGCAAGCCAGTG
MATR3KO #1
B
MATR3 (chr5, q31)
41 kb
ONT
HAP1 WT c.1
HAP1 WT c.7
MATR3KO c.1
MATR3KO c.5
Illumina
HAP1 WT c.1
HAP1 WT c.1
HAP1 WT c.7
HAP1 WT c.7
MATR3KO c.1
MATR3KO c.1
MATR3KO c.5
MATR3KO c.5
C
WT
MATR3KO
1
7
1
5
clone
MATR3
total
protein
D
MATR3
PTBP1
DAPI
HAP1 WT #1
MATR3KO #1

**Fig. S1. Deletion of *MATR3* in HAP1 cells.**

(**A**) *Top*, schematic depicts known protein domains within MATR3. Locations of four missense mutations associated with familial ALS are shown (*18*). NES, nuclear export sequence; ZF, zinc finger motif; RRM, RNA-recognition motif; NLS, nuclear localization sequence. *Middle*, Schematic depicts exon-intron structure of *MATR3* gene. The following gRNA GCCAAATACTAACGCTGGTG targeting exon 8 (denoted by arrow) was used for CRISPR-mediated genome editing. *Bottom*, the results of Sanger sequencing of deletions for MATR3 KO clones #1 and #5 are shown aligned to the corresponding WT reference sequence. (**B**) IGV screenshots depict normalized RNA-seq coverage tracks for the indicated samples either using ONT or Illumina. The window length and chromosomal location are indicated above the tracks, while the *MATR3* transcript direction (arrowheads) and exon-intron structure are indicated below the tracks based on annotated Reference Sequence (RefSeq) transcripts. (**C**) Western blot depicts abundance of MATR3 protein in two WT controls and two MATR3 KO clones. *Bottom*, an inset of the total protein stain is included as a loading control. (**D**) Immunofluorescence microscopy of a MATR3 KO clone compared to a WT control stained for MATR3 (red) and PTBP1 (green) proteins. Nuclei were stained with DAPI (blue). A representative image of each is shown. Scale bar is 10µm.

**Figure S2**

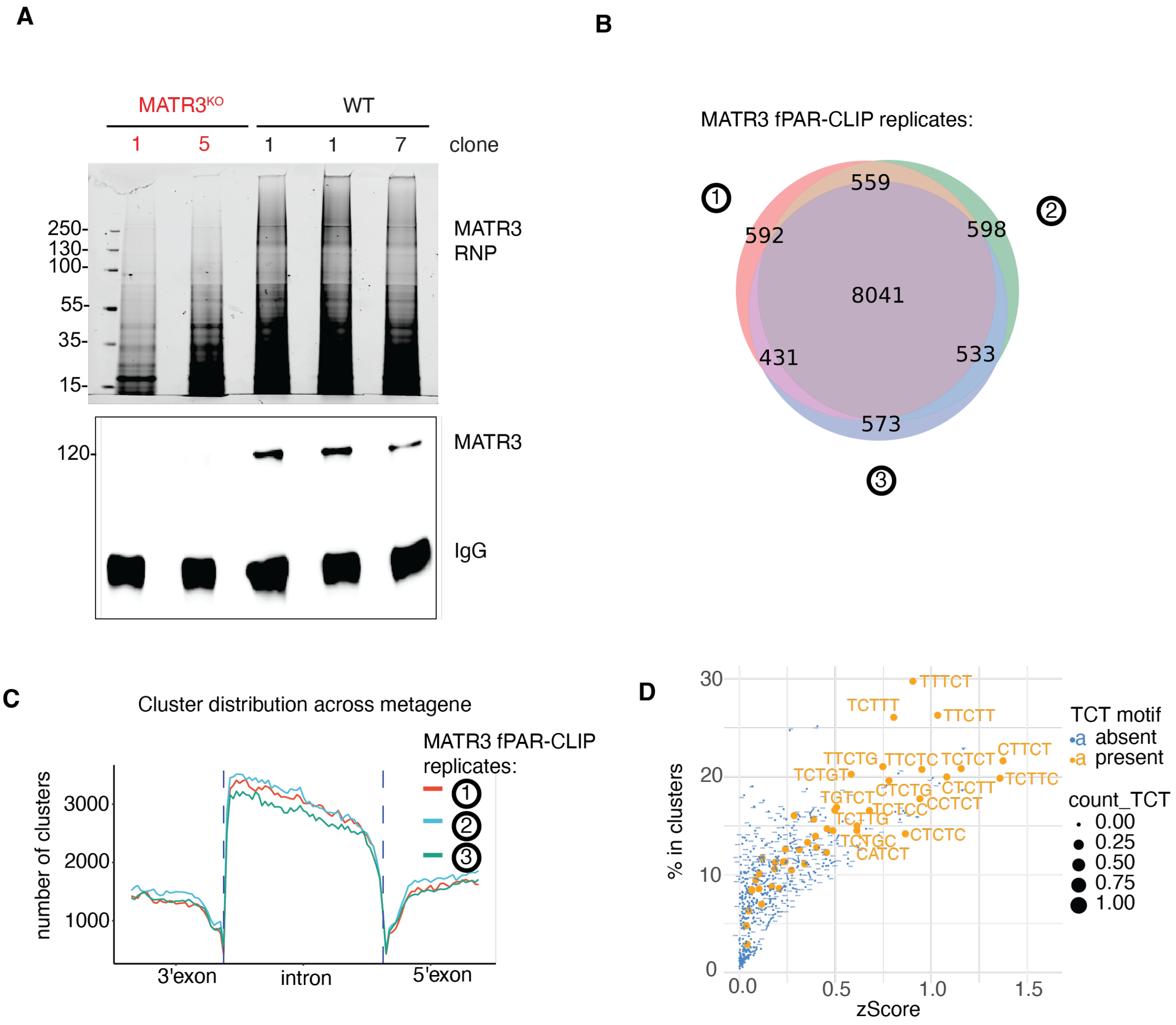

**Fig. S2. PAR-CLIP of endogenous MATR3.**

(**A**) Autoradiograph shows MATR3 crosslinked to $^{32}$P-labelled RNA fragments following immunoprecipitation and fractionation by SDS-PAGE. Two MATR3 KO samples are included as negative controls alongside the three WT replicates. *Bottom*, Western blot detection of MATR3 following PAR-CLIP is shown. (**B**) Venn diagram shows the overlap between three independent PAR-CLIP replicates. At least 8000 reproducible targets of endogenous MATR3 were identified. (**C**) Metagene analysis shows that MATR3 binding is enriched across introns for all three PAR-CLIP replicates. (**D**) Plot shows the 5-mer sequences containing TCT which are found to be enriched in MATR3 PAR-CLIPs. For comparison, not all motifs containing TCT are enriched. The motifs are pyrimidine-rich (C and U).

**Figure S3**

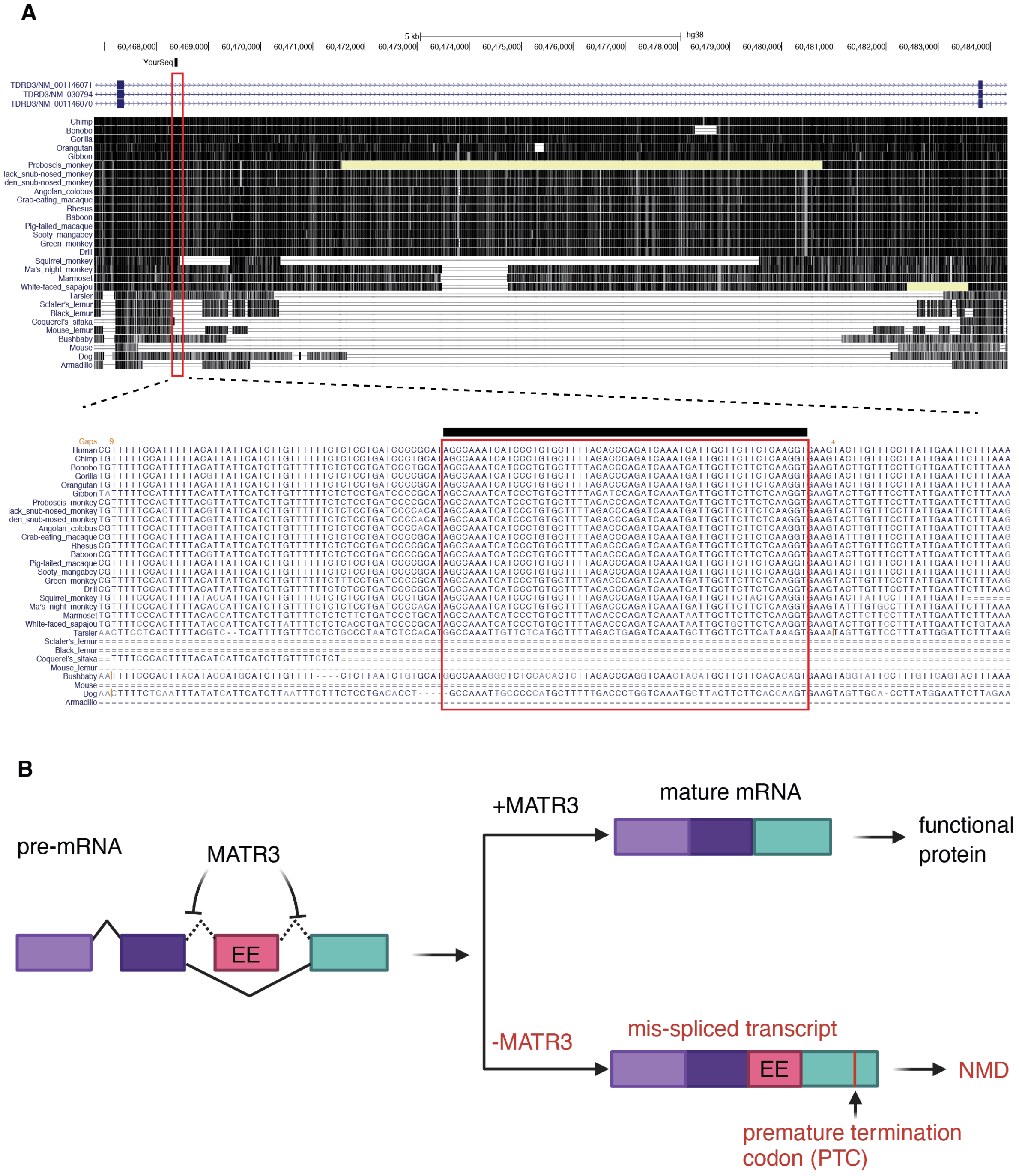
A
hg38
YourSeq
TDRD3/NM_001146071
TDRD3/NM_030794
TDRD3/NM_001146070
B
pre-mRNA
MATR3
EE
+MATR3
mature mRNA
functional protein
-MATR3
mis-spliced transcript
NMD
premature termination codon (PTC)

**Fig. S3. Evolutionarily conservation of novel exon in *TDRD3* and model of MATR3's role in splicing.**

(**A**) *Top panel*, UCSC Genome Browser screenshot depicts location of novel erroneous exon (red rectangle) downstream of *TDRD3* exon 5. Conservation tracks are included for 30 mammals (27 primates). *Bottom panel*, sequence alignment at the nucleotide level shows that the erroneous exon (53 base pairs) is perfectly conserved among 21 primate species including the di-nucleotide splice acceptor and donor motifs. (**B**) Cartoon depicts inhibition of erroneous exon (EE) inclusion by MATR3. When MATR3 is present, it safeguards canonical splicing of target transcript (eg. *TDRD3*) to generate mature mRNA and protein. In the absence of MATR3, mis-splicing generates an aberrant isoform with a premature termination codon which triggers non-sense mediated decay (NMD).

**Figure S4**

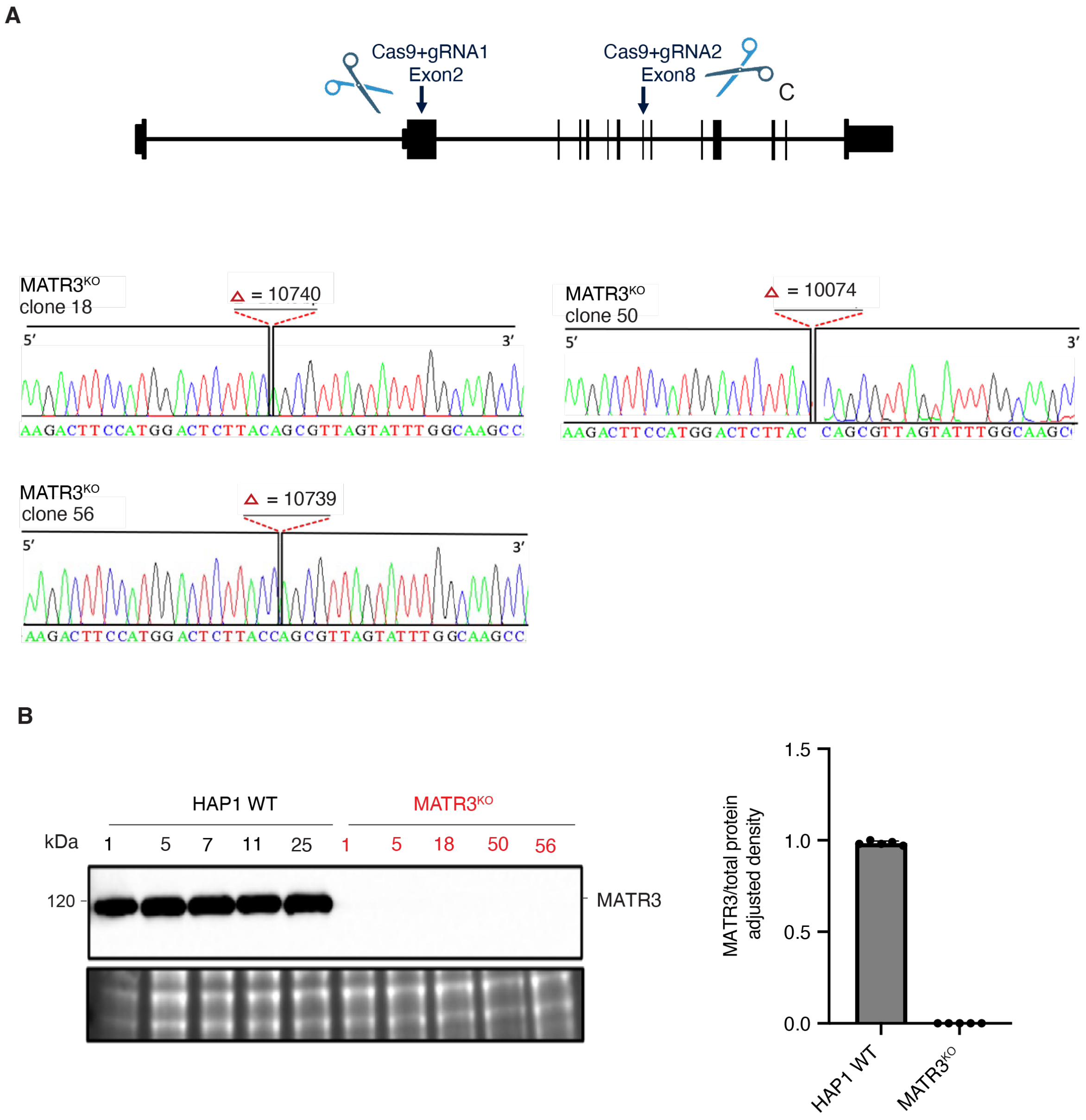

**Fig. S4. Additional deletions of *MATR3* in HAP1 cells.**

(**A**) *Top*, Schematic depicts exon-intron structure of *MATR3* gene. Two gRNAs targeting exons 2 and 8 (denoted by scissors) were used for CRISPR-mediated genome editing. *Bottom*, the results of Sanger sequencing of deletions for three additional MATR3 KO clones: #18, #50 and #56 are shown. Unique sizes of each deletion (Δ) are indicated. (**B**) Western blot depicts abundance of MATR3 protein in five WT controls and five MATR3 KO clones. Three new WT clones (#7, 11, and 25) were included as matching controls. An inset of the total protein stain is included as a loading control.

**Figure S5**

**A**

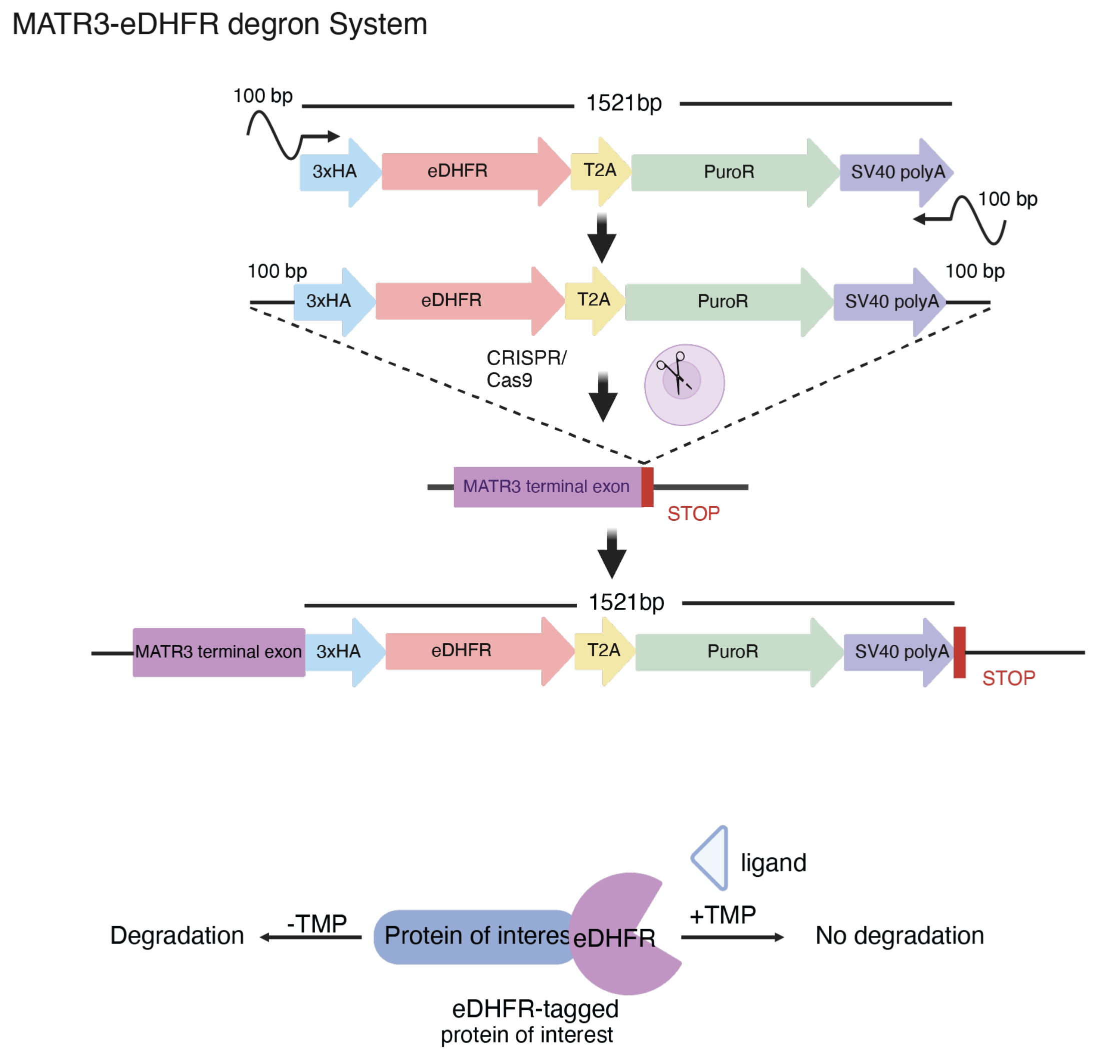

MATR3-eDHFR degron System
100 bp
1521bp
3xHA
eDHFR
T2A
PuroR
SV40 polyA
100 bp
100 bp
3xHA
eDHFR
T2A
PuroR
SV40 polyA
100 bp
CRISPR/
Cas9
MATR3 terminal exon
STOP
1521bp
MATR3 terminal exon
3xHA
eDHFR
T2A
PuroR
SV40 polyA
STOP
ligand
Degradation
-TMP
Protein of interes
eDHFR
+TMP
No degradation
eDHFR-tagged
protein of interest

**B**

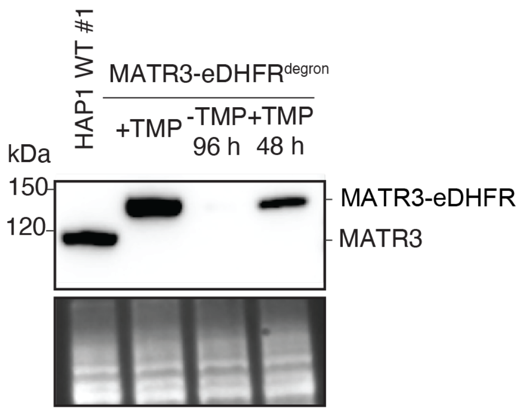

HAP1 WT #1
MATR3-eDHFR$^{degron}$
+TMP
-TMP
96 h
+TMP
48 h
kDa
150
120
MATR3-eDHFR
MATR3

**Fig. S5. MATR3-eDHFR degron and *TDRD3* mis-splicing in HCT116 cells after RNA interference of *MATR3*.**

(**A**) Top, schematic depicts the 1.5 kb HDR donor inserted in place of STOP codon in last exon of *MATR3* via CRISPR/Cas9. As a result, the C-terminus of MATR3 is fused to three tandem copies of the HA epitope tag (3xHA), followed by the eDHFR degron tag, a Thosea asigna virus 2A (T2A) sequence to mediate ribosomal skipping. The puromycin-resistance cassette is for drug selection of stable chromosomal integration events. The small molecule trimethoprim (TMP) binds to eDHFR and stabilizes the fusion protein. When TMP is removed, the eDHFR fusion protein is degraded. (**B**) Western blot of MATR3 (*top*) is shown. HAP1 cells engineered to express endogenous MATR3 that is eDHFR-tagged were grown in 10 μM trimethoprim (TMP) to stabilize MATR3-eDHFR fusion protein (lane 2); then washed and grown in TMP free (-TMP) media for 96 h to deplete MATR3-eDHFR (lane 3); and then in 10 μM TMP for 48 h to restore MATR3-eDHFR expression (lane 4). *Bottom*, an inset of the total protein stain is included as a loading control.

### Figure S6

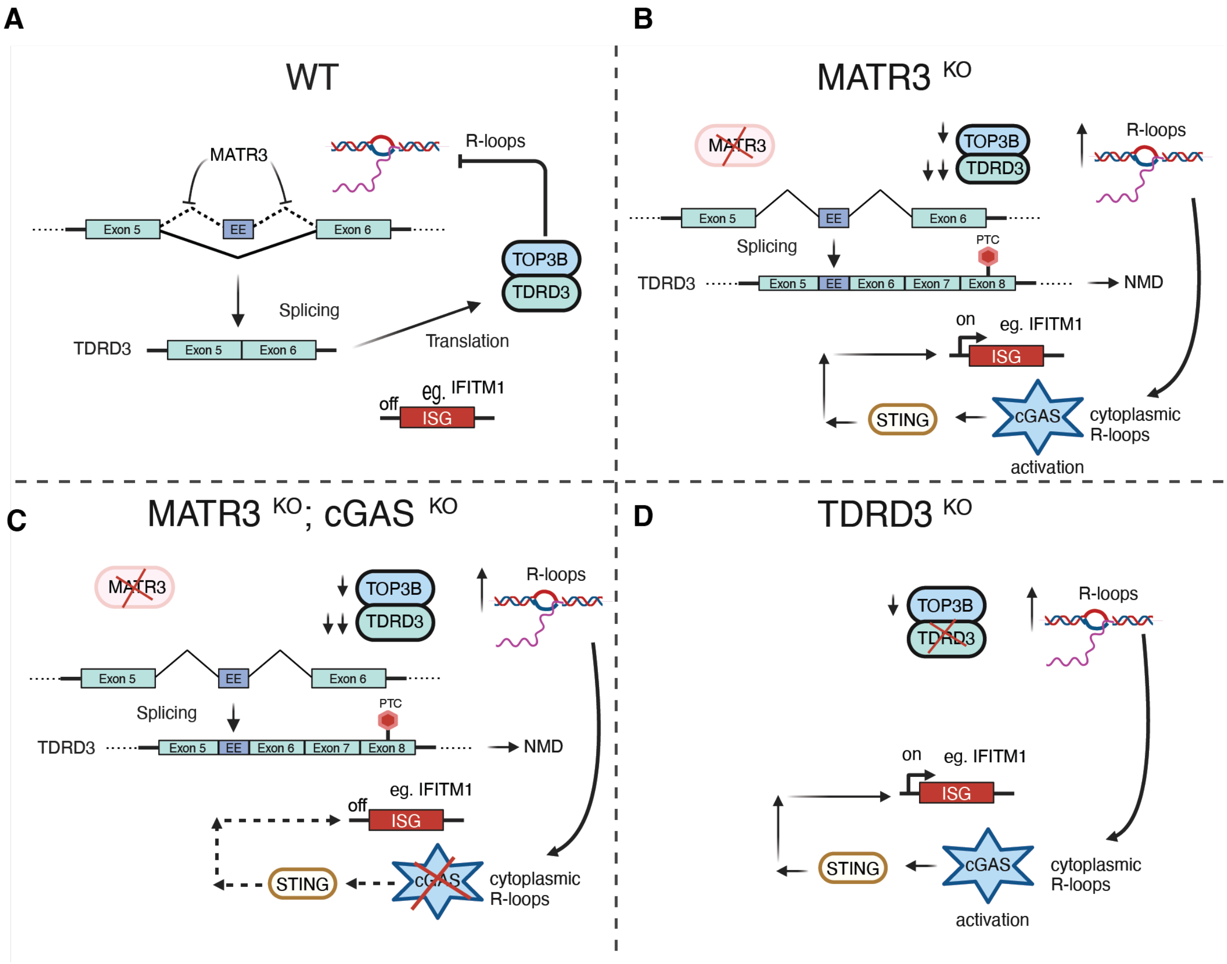

**Fig. S6. Proposed working model.**

(**A**) In WT cells, MATR3 ensures canonical splicing of TDRD3. The TDRD3-TOP3B complex maintains homeostasis of R-loops to levels which are tolerated. ISGs are turned off. (**B**) In MATR3 KO cells, mis-splicing of TDRD3 results in a PTC and NMD. As a result, TDRD3 protein is diminished, and TOP3B is destabilized. Aberrant accumulation of R-loops lead to spillover of RNA-DNA hybrids to the cytoplasm as previously described (*19*). Cytoplasmic RNA-DNA hybrids are sensed by cGAS and activates STING which lead to induction of ISGs similar to what happens in RNaseH2-deficient cases of AGS (*20*). (**C**) The interferonopathy-like phenotype in MATR3 KO cells can be treated by targeting cGAS. (**D**) The interferonopathy in MATR3 KO cells is phenocopied by TDRD3 KO.

**Table S1.**
List of oligonucleotides.

| Sequence (5' to 3') | Name | Supplier |
|---|---|---|
| CRISPR gRNA | | |
| CTTCCATGGACTCTTACCGA | MATR3 exon 2 | IDT |
| GCCAAATACTAACGCTGGTGG | MATR3 exon 8 | IDT |
| AAGGAAACTTAAGATGTGCA | MATR3 exon 14 | IDT |
| GGGCCGAACTTTCCCGCCTT | CGAS exon 1 | IDT |
| AGACTCGGTGGGATCCATCG | CGAS exon 5 | IDT |
| TCGCAATGTTGCTGCACCAA | TDRD3 exon 4 | IDT |
| GTGTGTATCTCATGTCCAAG | TDRD3 exon 7 | IDT |
| AGAAGCATACTATCTGTTAC | TDRD3 intron 6 upstream | IDT |
| GATTTGGCTATGCGGGGATC | TDRD3 intron 6 downstream | IDT |
| Genotyping primers | | |
| AGATGGAGAAAGGTGTAGGGA | MATR3 exon 2 forward | IDT |
| TTAGCAATAATTTGAAATAACCACAGTT | MATR3 exon 8 reverse | IDT |
| TTCCCCGTGCCTTATTGTAATG | MATR3 exon 14 forward | IDT |
| GGAAGTCAATCTCTACAAAGTACTTCTGTC | MATR3 exon 14 reverse | IDT |
| CACAGCCCTGAATAGAGTTTCAT | TDRD3 intron 6 forward | IDT |
| TTCAGCAATAGCAGCCGTCC | TDRD3 intron 6 reverse | IDT |
| RT-PCR primers | | |
| CTGAACACACCACCTGGAACTA | TDRD3 exon 5 forward | IDT |
| GCAATCATTTGATCTGGGTC | TDRD3 EE reverse | IDT |
| CCAAATCATCCCTGTGCTTT | TDRD3 EE forward | IDT |
| CTGTCCAAAAGGCACAAAAG | TDRD3 exon 6 reverse | IDT |
| AGGATTTGGAAAGGGTGTTTATTC | HPRT1 exon 2 forward | IDT |
| CAGAGGGCTACAATGTGATGG | HPRT1 exon 3 reverse | IDT |
| HDR donor primers | | |
| T*T*GCTGCATTTCTCTTAGGTGACTTAATGGCTGTA ATTCTCTTTCTTTATAGAAATTTCTGAATAAATTGGCA GAAGAACGCAGACAGAAGAAGGAAACTGGAGGCGG TTACCCATAC | MATR3_3xHA forward | IDT |
| G*T*TTTCTTCTAACTATCAGTATTGTATTTAAAAAAG GTTAACATTAAAAACAAAGAACCATTATTTTCTTTGA AATCATTAAATCTCCTTGCACATCTTAA GTCGACTGATCATAATCAGC | SV40_MATR3 reverse | IDT |

* phosphorothioate linkages

**Table S2.**
List of antibodies.

| Antibody | Supplier | Catalog# |
|---|---|---|
| Rabbit polyclonal anti-MATR3 (used for immunostaining) | Sigma | HPA036565 |
| Rabbit polyclonal anti-MATR3 (used for PAR-CLIPs) | Bethyl | A300-591A |
| Rabbit monoclonal anti-MATR3 (used for Westerns) | ThermoFisher | MA5-34642 |
| Rabbit polyclonal anti-PTBP1 | ThermoFisher | 32-4800 |
| Rabbit monoclonal anti-TDRD3 | CST | 5942S |
| Rabbit polyclonal anti-TOP3B | Abcam | ab183520 |
| Mouse monoclonal anti-DNA-RNA hybrids (clone S9.6) | Sigma | MABE1095 |
| Rabbit polyclonal anti-IFITM1 | Proteintech | 60074-1-Ig |
| Rabbit monoclonal anti-cGAS | CST | 79978s |
| Rabbit monoclonal anti-MARCKS | CST | 5607s |
| Rabbit polyclonal anti-PSMB8 | ThermoFisher | PA1-972 |
| HRP-conjugated anti-rabbit IgG 2° antibody | CST | 7074S |
| HRP-conjugated anti-rabbit IgG 2° antibody | CST | 7076S |
| Alexa 488 conjugated anti-rabbit IgG 2° | ThermoFisher | A27034 |
| Alexa 488 conjugated anti-mouse IgG 2° | ThermoFisher | A28175 |
| Alexa 594 conjugated anti-rabbit IgG 2° | ThermoFisher | A11012 |

**Movie S1.**
Animated 3-D reconstruction of S9.6 immunostaining to demonstrate the cytoplasmic localization of RNA-DNA hybrids.

**Data S1. (separate file)**
Results of DESeq2 analysis of Illumina RNA-seq data comparing MATR3 KO versus WT.

**Data S2. (separate file)**
Results of DESeq2 analysis of ONT RNA-seq data comparing MATR3 KO versus WT.

**Data S3. (separate file)**
Results of nSolver analysis of Nanostring data comparing MATR3 KO versus WT.

**Data S4. (separate file)**
List of clusters (target sites) identified from MATR3 PAR-CLIP. Data from three replicates are on separate sheets within.

**Data S5. (separate file)**
Merged results of three MATR3 PAR-CLIP replicates above.

**Data S6. (separate file)**
Summary of rMATS analysis of Illumina RNA-seq data comparing MATR3 KO versus WT.

**Data S7. (separate file)**
Prioritized short list of targets.
1) ~8000 PAR-CLIP targets were intersected with ~800 SE genes from rMATS
2) target list was ranked based on fold-change (MATR3 KO vs WT). The top 47 are shown.
